# Propagation of cortical activity via open-loop intrathalamic architectures: a computational analysis

**DOI:** 10.1101/574178

**Authors:** Jeffrey W. Brown, Aynaz Taheri, Robert V. Kenyon, Tanya Berger-Wolf, Daniel A. Llano

**Affiliations:** University of Illinois College of Medicine at Urbana-Champaign, Urbana, IL 61801; Department of Computer Science, University of Illinois at Chicago, Chicago, IL 60607; Beckman Institute for Advanced Science and Technology, University of Illinois at Urbana-Champaign, Urbana, IL 61801; Neuroscience Program, University of Illinois at Urbana-Champaign, Urbana, IL 61801; Department of Molecular and Integrative Physiology, University of Illinois at Urbana-Champaign, Urbana, IL 61801

**Keywords:** Thalamus, thalamic reticular nucleus, intrathalamic signaling, cortical signaling, open-loop, propagation, computational model

## Abstract

Propagation of signals across the cerebral cortex is a core component of many cognitive processes and is generally thought to be mediated by direct intracortical connectivity. The thalamus, by contrast, is considered to be devoid of internal connections and organized as a collection of parallel inputs to the cortex. Here, we provide evidence that “open-loop” intrathalamic connections involving the thalamic reticular nucleus (TRN) can support propagation of oscillatory activity across the cortex. Recent studies support the existence of open-loop thalamo-reticulo-thalamic (TC-TRN-TC) synaptic motifs in addition to traditional closed-loop architectures. We hypothesized that open-loop structural modules, when connected in series, might underlie thalamic and, therefore cortical, signal propagation. Using a supercomputing platform to simulate thousands of permutations of a thalamo-reticular-cortical network and allowing select synapses to vary both by class and individually, we evaluated the relative capacities of closed- and open-loop TC-TRN-TC synaptic configurations to support both propagation and oscillation. We observed that 1) signal propagation was best supported in networks possessing strong open-loop TC-TRN-TC connectivity; 2) intrareticular synapses were neither primary substrates of propagation nor oscillation; and 3) heterogeneous synaptic networks supported more robust propagation of oscillation than their homogeneous counterparts. These findings suggest that open-loop heterogeneous intrathalamic architectures complement direct intracortical connectivity to facilitate cortical signal propagation.

**Significance Statement:** Interactions between the dorsal thalamus and thalamic reticular nucleus (TRN) are speculated to contribute to phenomena such as arousal, attention, sleep, and seizures. Despite the importance of the TRN, the synaptic microarchitectures forming the basis for dorsal thalamus-TRN interactions are not fully understood. The computational neural model we present incorporates “open-loop” thalamo-reticular-thalamic (TC-TRN-TC) synaptic motifs, which have been experimentally observed. We elucidate how open-loop motifs possess the capacity to shape the propagative properties of signals intrinsic to the thalamus and evaluate the wave dynamics they support relative to closed-loop TC-TRN-TC pathways and intrareticular synaptic connections. Our model also generates predictions regarding how different spatial distributions of reticulothalamic and intrareticular synapses affect these signaling properties.

## Introduction

Propagation of activity across the cerebral cortex is thought to underlie multiple cognitive processes, as well as pathological processes such as epilepsy and migraine (1–4). Cortical regions are highly interconnected via direct axonal projections as well as via polysynaptic pathways involving the basal ganglia and thalamus (5, 6). Cortical signal propagation is generally thought to be mediated via direct cortical connections (7, 8), but recent evidence suggests that the thalamus serves as a control point to modify cortical activity during cognitive processes such as attentional shifting (9). An advantage of a thalamic mode of signal propagation is the efficiency by which modulatory influences may control thalamic, and therefore cortical, propagation. The thalamus, however, is generally thought to have limited internal connectivity and therefore limited capacity to serve as a substrate for signal propagation.

A major intermediary allowing for communication between thalamocortical neurons, the thalamic reticular nucleus (TRN), is a sheet of GABAergic neurons that partially envelops the dorsal thalamus (10). It has been speculated to participate in phenomena ranging from selective attention (11–13) to sleep and arousal (12–15) and fear responses (16), and may play a role in generating absence seizures (17–21), symptoms of neurodevelopmental disorders (22, 23), and schizophrenia (24). The TRN projects exclusively to TC neurons, while receiving reciprocal, glutamatergic thalamoreticular (TC-TRN) connections (25).

The structural microarchitecture of bidirectional pathways connecting the dorsal thalamus and TRN has been the subject of ongoing debate. It was originally assumed that thalamo-reticulo-thalamic (TC-TRN-TC) pathways were reciprocal, forming “closed loops” of recurrent inhibition delivered to TC neurons (Fig. 1A, left) (10, 15, 26–28). While closed disynaptic loops have indeed been confirmed, they were only identified in a minority of examined TC-TRN pairs (10, 29–33). Another connectional scheme between the dorsal thalamus and TRN is the so-called “open-loop” TC-TRN-TC pathway, wherein a TC neuron is not reciprocally inhibited by the TRN neuron it excites (Fig. 1A, right). Open-loop configurations have been inferred from recordings in rodent thalamic slice preparations (34–38) and confirmed in anatomical studies (32, 39, 40). Furthermore, open-loop pathway variants in the form of X-TRN-TC are also known to exist, with X representing indirect sources of modulation to the sensory thalamus via the TRN, such as monoaminergic and cholinergic brainstem nuclei, nuclei of the basal forebrain, amygdala, and prefrontal cortex (9, 41–45).

**Figure 1.**
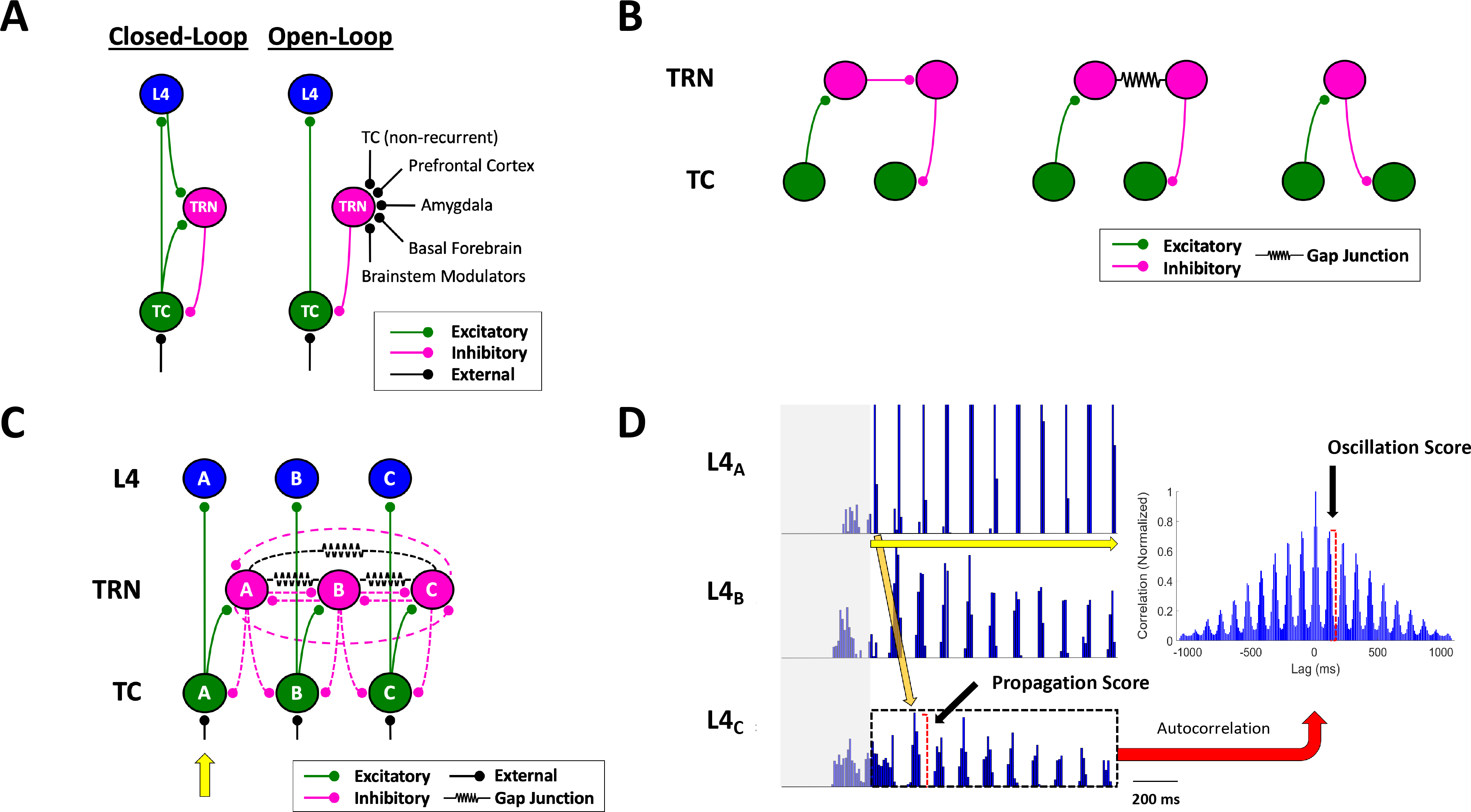
Pathways and properties of thalamocortical signaling. **A:** Closed- vs. open-loop thalamo-reticulo-thalamic configurations. **B:** Three possible pathways through which a signal might propagate from one thalamocortical (TC) neuron to another via the thalamic reticular nucleus (TRN). **C:** Baseline thalamo-reticulo-cortical model network. Broken-line synapses were allowed to vary either as a class (homogeneously) or independently of one another (heterogeneously). **D:** Sample L4 spike histograms (detrended) in a network permutation responding to a fixed, sustained stimulus delivered to TC_A_ (yellow arrow). The propagation score assigned to any network permutation was quantified as the amplitude of the initial stimulus-evoked response in the detrended L4_c_ histogram; response propagation across the L4 subnetwork (orange arrow) was consistently linear, and thus the initial response in L4_C_ was observed at a fixed interval relative to the onset of stimulation. Oscillation intrinsic to any network variant was quantified as the amplitude of the first off-center peak in the normalized autocorrelogram (right) of post-stimulation activity in the detrended L4_C_ histogram (within broken black box). The initial 400 ms of activity preceding the fixed stimulus (in grey) is shown here for each histogram but was not included in the calculations of either propagation or oscillation. Note that the bin heights in the L4_A_ histogram shown here were truncated in order to maintain identical vertical scaling across all three L4 histograms.

We previously observed through a computational model that the open-loop TC-TRN-TC pathway, rather than uniformly depressing thalamic (and consequently cortical) activity, paradoxically enhanced thalamocortical output over a range of TC and TRN input frequencies (46). This finding demonstrated the capacity of an open-loop system to function as a tunable filter of thalamocortical transmission, subject to the temporal dynamics of input to the TRN, whether from other, non-reciprocally connected TC neurons or extrinsic sources. In both our previous model and earlier models built on closed-loop TC-TRN-TC synaptic motifs, the post-inhibitory rebound exhibited by TC neurons, as mediated by T-type Ca^2+^ channels and driven by inhibition from the TRN, served as a catalyst of signal propagation within the networks (9, 46–55).

Based on previous studies of open-loop TC-TRN-TC synaptic organization, we hypothesized that these synaptic modules might underlie intrathalamic and therefore intracortical signal propagation. Accordingly, we sought here to evaluate the efficacy of open-loop pathways relative to other potential synaptic configurations in mediating signal transmission across the thalamus and cortex. To this end, we constructed a model network based on that of (46) by connecting in series three thalamo-reticulo-layer-4-cortical (TC-TRN-L4) pathways, potentially featuring both closed- and/or open-loop TC-TRN-TC motifs, with the latter constituting one mode of connectivity between parallel TC-TRN-L4 pathways. Intrareticular synapses represented the other structural connections between pathways, based on the identification of both GABAergic (56–62) and electrical synapses (61–65) between TRN neurons. Thus, we included three different polysynaptic configurations between vertical pathways in our network (Fig. 1B, from left to right): 1) those with a chemical intrareticular synapse; 2) those with anelectrical intrareticular synapse; and 3) open-loop TC-TRN-TC pathways. To analyze how each variety of inter-pathway connection contributed to network dynamics, permutations of the baseline network were generated by varying three properties associated with each of the inter-pathway synaptic motifs. We quantified network dynamics as a function of variable TC-TRN-TC and intrareticular synaptic architectures by defining and measuring two properties inherent to stimulus-evoked responses in each network variant: propagation and oscillation, with the latter included in light of the fact that many characterized thalamic waveforms both oscillate and propagate through the thalamus and cortex (25).

## Network architecture and simulations

We constructed a neuronal network comprising three interconnected thalamo-reticulo-cortical pathways (Fig. 1C). Thalamic, reticular, and cortical cell layers were aligned topographically, such that TC_A_ projected to both TRN_A_ and L4_A_ (10, 25, 52, 66, 67).

In the case of homogeneously varied synaptic network permutations, the synaptic parameters associated with three inter-pathway motifs varied as a class, with all external, TC-TRN, and TC-L4 synaptic conductances held constant: 1) GABAergic intrareticular (TRN-TRN_GABA_) synapses ranged in conductance between 0 and 450 nS; 2) electrical intrareticular (TRN-TRN_Elec_) synapses ranged in coupling coefficient between 0 and 0.36; and 3) a TC-TRN-TC “openness” coefficient, defined as the weight distribution of lateral (open-loop, comprising 2 synapses of the form TRN_*i*_➔TC_*i*+*1*_) vs. recurrent (closed-loop, comprising 3 synapses of the form TRN_*i*_➔TC_*i*_) reticulothalamic connectivity, varied between 0 (completely closed-loop) and 1.0 (completely open-loop) and with a baseline TRN-TC conductance of 80 nS.

For the heterogeneously varied synaptic network variants, all TRN-TRN and TRN-TC synapses were allowed to vary independently. Domains for each of the synaptic variables were selected to include the range of conductance or coupling strengths reported in physiological measurements and/or used in similar neural models (19, 49, 52, 54, 63, 64, 67).

Ongoing afferent synaptic input was delivered to every TC neuron in the model as Poisson-modulated spike trains centered at 40 Hz. An additional 200-Hz pulse train was applied to neuron TC_A_ between *t*=0.400 and *t*=1.500 s during every network simulation run. This high-frequency stimulus was modeled on those used to elicit spindle-like waves in a ferret thalamoreticular slice preparation (18, 68). A given network’s output was compiled by assembling spike histograms (10-ms bins) averaging 1,000 simulations for every L4 neuron (Fig. 1D). Network properties were quantified in the most downstream element of the cortical output layer, L4_C_. Propagation across a network was quantified as the amplitude of the initial stimulus-evoked response in the detrended L4_C_ histogram. The degree of oscillation supported by each network permutation was defined as the amplitude of the first off-center peak in the normalized autocorrelogram of post-stimulation activity (Fig. 1D). Both propagation and oscillation scores are reported as normalized to the maximum scores tabulated for each property. Given the high prevalence of propagating oscillatory waves in the cerebral cortex [reviewed in (69)], we furthermore defined a composite “optimization” (*Op*) metric to measure the capacity of networks to simultaneously support and balance between propagation (*Pr*) and oscillation (*Os*):

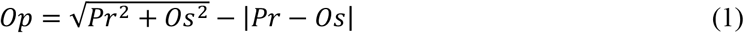

## Results

### Homogeneously varied synaptic models

Stimulus-evoked responses propagated linearly across the length of homogeneous synaptic networks, occurring at average fixed intervals of 93.31 ± 0.35 ms (mean ± standard error of the mean; range, 60-110 ms) between adjacent TC-TRN-L4 pathways, across all model permutations and with a mean velocity of 0.54 mm/s, assuming a 50 µm separation between adjacent neurons in each network layer. All 770 homogeneous network variants were ranked according to their cortical propagation scores (Fig. 2A, top). Linear regression analysis (*R*^*2*^=0.793, root-mean-square-error or RMSE=0.047, *p*<0.0001) demonstrated a strong positive correlation between the TC-TRN-TC openness coefficient and propagation score (normalized regression coefficient or NRC=1.000). By contrast, chemical and electrical TRN-TRN synaptic connectivity tended to modestly diminish propagation (NRC=−0.173 and NRC=−0.136, respectively; Table S1). Further, other excitatory connectivity, such as cortico-cortical or corticothalamic connectivity, often postulated as being important for cortical signal propagation (5, 7, 8), was not necessary. Thus, the homogeneously varied synaptic network permutations that best accommodated signal propagation were generally ones with weak or absent synapses between TRN neurons and strong open-loop TC-TRN-TC connections. For example, Network a, which epitomizes this architecture, exhibited robust signal propagation in response to a fixed stimulus delivered to TC_A_; a representative simulation of this network is shown in Fig. 2B, left, and its position in Fig. 2C is labeled. Stimulus-evoked activity in this network tended to propagate efficiently from L4_A_ to L4_C_: near-synchronous propagation cascades were elicited in both the TRN and L4 layers of the model, having been stimulated by propagating activity in upstream TC neurons. Smooth, linear propagation of action potentials across the network depended on the synchronous induction of inhibitory postsynaptic potentials (IPSPs) and the ensuing post-inhibitory rebound spikes in TC neurons, which occurred reliably and at fixed intervals in Network a.

**Figure 2.**
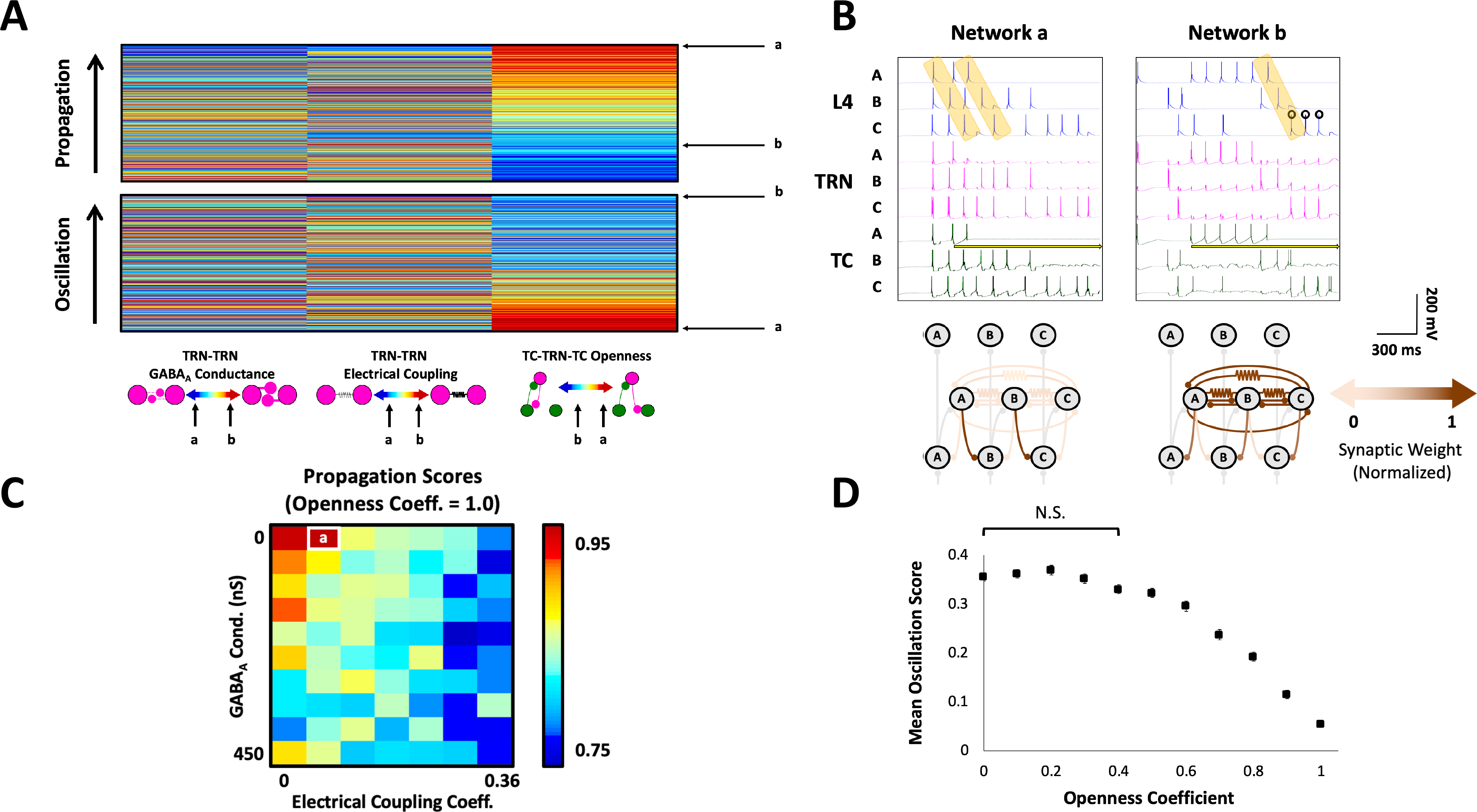
Propagation and oscillation in homogeneously varied synaptic networks (N=770). **A:** Ordinal heat maps ranking homogeneously varied synaptic network permutations according to the extent of supported signal propagation and oscillation. The network property ranks and synaptic makeups of two selected networks, Networks a and b, are indicated. **B:** Representative simulations and circuit diagrams depicting the normalized synaptic makeups for the two selected networks. The yellow arrow indicates when the fixed stimulus was delivered to TC_A_ in each simulation. Orange highlighting indicates epochs of linear propagation, while circles are placed above spikes occurring during periods of oscillatory activity. **C:** A heat map displaying propagation scores in TRN-TRN synaptic parameter space for the 70 fully open-loop networks (openness coefficient=1.0), with Network a highlighted. **D:** Mean oscillation scores for networks varied nonlinearly as a function of their openness coefficients, with networks possessing openness coefficients of 0 and 0.4 supporting oscillation to equal extents (one-way ANOVA with Tukey post-hoc tests, *F*(10,759)=137.8, *p*<0.0001). Error bars indicate standard errors of the mean; N.S.=not significant.

A 2° multiple regression model of propagation as a function of all three synaptic class variables (*R*^*2*^=0.842, RMSE=0.041, *p*<0.0001; Table S1) revealed modestly negative interaction term between TRN-TRN_Elec_ synapses and TC-TRN-TC openness (NRC=−0.365), indicating that in networks where both electrical synapses were strong and TC-TRN-TC openness high, the extent of supported propagation diminished nonlinearly; a smaller negative interaction between TRN-TRN_GABA_ synapses and TC-TRN-TC openness was also observed (NRC=−0.152). Together, these terms suggested that propagation was more significantly affected by connections in the TRN layer as a function of increasing open-loop TC-TRN-TC architecture. This relationship is evident in Fig. 2C, as propagation scores conspicuously decreased in network variants with an openness coefficient of 1.0 as either chemical or electrical synapses increase in weight.

Oscillatory responses recurred in L4_C_ neurons at a mean frequency of 9.07 ± 0.2 Hz (range, 7.14-12.50 Hz) across all homogeneous model permutations. Propagation and oscillation scores across all 770 homogeneous networks were strongly anticorrelated (Pearson’s *r*=−0.671, *p*<0.0001). Accordingly, oscillation was best accommodated in network permutations exhibiting strongly closed-loop connectivity (Fig. 2A, bottom), however the capacity to support oscillation was neither markedly linear nor monotonically decreasing as a function of increasing openness coefficient (Fig. 2D). Rather, a one-way analysis of variance (ANOVA) with Tukey’s tests revealed that, on average, oscillation scores peaked and remained statistically indistinguishable from one another across the subset of network permutations with openness coefficients between 0 and 0.4, with scores then decreasing in a roughly linear fashion with increasing TC-TRN-TC openness [*F*(10,759)=137.8, *p*<0.0001]. These data suggest that networks with mixed open- and closed-loop connectivity (which is likely close to physiological reality) can support the coexistence of oscillation and propagation (see *Heterogeneously varied synaptic models*, below).

The predominant mechanism by which oscillation arose in L4_C_ was through post-inhibitory rebound in TC_C_, as engendered by the strong recurrent inhibition found in network permutations exhibiting primarily closed-loop TC-TRN-TC connectivity. This mode of oscillation was exemplified by Network b, a strongly closed-loop network variant. In the simulation shown of this network (Fig. 2B, right), oscillatory activity was enabled by a single epoch of signal propagation. Notably, neither the presence of strong GABAergic nor electrical intrareticular synapses in Network b exerted much effect on its ability to support oscillation, as predicted by the regression models.

### Heterogeneously varied synaptic models

Recent studies have highlighted heterogeneity in TRN neuronal connectivity, synaptic physiology and chemical identities (70–72). We therefore examined the impact of allowing all synaptic connections involving the TRN to be independently varied. We constructed circuit-level schematics of linear regression models for propagation (Fig. 3A, top) and oscillation (Fig. 3A, bottom) as functions of the 14 synaptic variables in heterogeneous networks.

**Figure 3.**
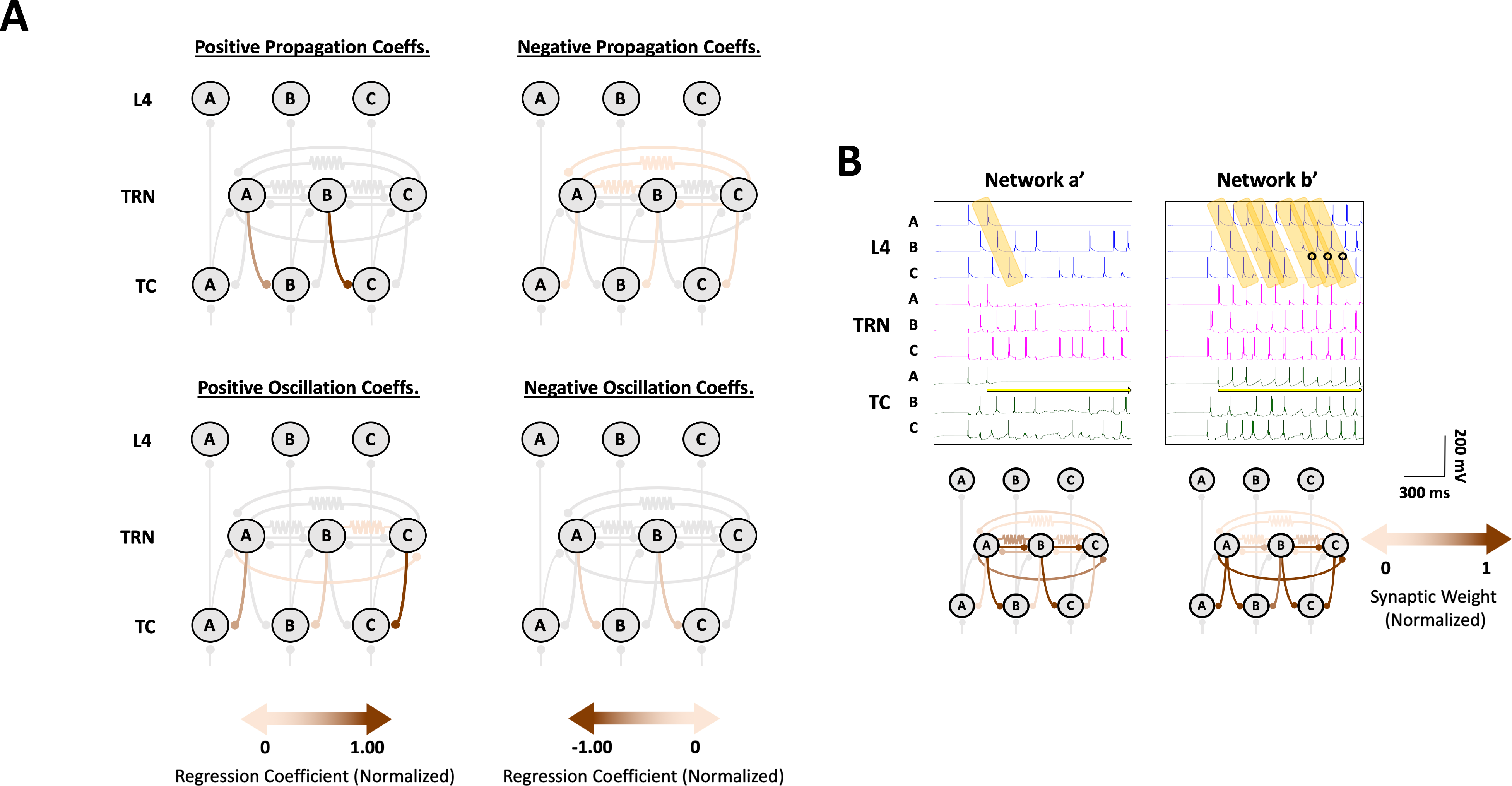
Propagation and oscillation in heterogeneously varied synaptic networks (N=12,681). **A:** Network regression models illustrating how propagation (top) and oscillation (bottom) varied as a function of individual synaptic weights across simulated heterogeneously synaptic network permutations. Gray synapses are either non-variable or associated with normalized regression coefficients with absolute values under 0.05. Synapses with positive and negative coefficients in the regression models are depicted separately in the left- and right-sided circuit diagrams, respectively. **B:** Representative simulations for two selected heterogeneous networks, whose normalized synaptic weights are depicted in the circuit diagrams. Networks a’ and b’ respectively illustrate propagation and propagation of oscillation from Column A to Column C.

Propagation in heterogeneously varied synaptic networks increased chiefly as a function of increasing the strength of the more downstream of the two laterally inhibitory TRN-TC synapses, TRN_B_➔TC_C_: the corresponding term in a linear regression model of propagation (*R*^*2*^=0.742, RMSE=0.069, *p*<0.0001; Table S2) possessed an NRC of 1.000 (Fig. 3A, top). Propagation scores also scaled to a lesser extent with the more upstream laterally inhibitory reticulothalamic synapse, TRN_A_➔TC_B_ (NRC=0.608). The two inhibitory intrareticular synapses originating at the rightmost end of the model network, TRN_C_➔TRN_A_ and TRN_C_➔TRN_B_, both exerted a small negative effect on propagation (NRC=−0.087 and NRC=−0.084, respectively). Additionally, two TRN-TRN_Elec_ synapses, TRN_A_=TRN_B_ and TRN_A_=TRN_C_ (where the “=” denotes an electrical synapses), marginally decremented propagation in heterogeneous networks, with NRCs of −0.051 and −0.072, respectively. These findings at an individual synaptic level comported with the observation that strong TRN-TRN interactions, whether chemical or electrical, tended to impede signal propagation in homogeneous network variants.

A 2° regression model (*R*^*2*^=0.857, RMSE=0.051, *p*<0.0001; Table S2) disclosed a large, propagation-enhancing interaction between the two laterally inhibitory synapses (NRC=0.753), underscoring the same dependence of propagation on strong open-loop TC-TRN-TC connectivity as seen in homogeneously synaptic networks, but additionally demonstrating that propagation scores increased nonlinearly as a function of simultaneously increasing the weights of TRN_A_➔TC_B_ and TRN_B_➔TC_C_. Interactions between TRN-TRN synapses of either variety and TRN-TC synapses tended diminish propagation, as did those between recurrent and lateral inhibitory TRN-TC synapses. Taken together, the linear and 2° regression models indicated that heterogeneous network permutations with strong laterally inhibitory TRN-TC synapses tended to best support propagation. Consistent response propagation across the length of the network was epitomized by Network a’, in which TRN_A_➔TC_B_ and TRN_B_➔TC_C_ were both relatively strong and those synapses impeding propagation relatively weak (Fig. 3B, left).

### Comparisons between homogeneously and heterogeneously varied synaptic architectures

In contrast to the homogeneous models, there was a very small negative correlation between the propagation and oscillation scores of these networks (*r*=−0.0296, *p*=0.0008), suggesting that propagation and oscillation more easily coexist in heterogeneous than homogeneous models. This supposition was confirmed through a 2° regression analysis (*R*^*2*^=0.388, RMSE=0.118, *p*<0.0001), which suggested that interactions between recurrently and laterally inhibitory TRN-TC synapses (NRCs ranging between 0.345 and 0.669) facilitated the propagation of oscillation, a mechanism typified by Network b’ (Fig. 3B, right). Two intrareticular synapses, TRN_A_-TRN_C_ and TRN_A_=TRN_C_, tended to contribute modestly to oscillation (NRCs of 0.115 and 0.117, respectively, in the linear regression model, *R*^*2*^=0.253, RMSE=0.131, *p*<0.0001; Fig. 3A, bottom), while, in their individual capacities, TRN_A_➔TC_B_ and TRN_B_➔TC_C_ diminished oscillation (NRCs of −1.000 and −0.892, respectively).

We analyzed the relative capacities of homogeneously and heterogeneously varied synaptic networks to support propagation, oscillation, and optimization by comparing the 20 highest scores achieved by homogeneous and heterogeneous network permutations with respect to each performance metric. No significant differences in mean propagation scores between top-performing homogeneous and heterogeneous networks were disclosed [unpaired *t*-test, *t*(38)=0.46, *p*=0.647; Fig. 4]. We attributed this lack of differences to the fact that network permutations in which the synapses TRN_A_➔TC_B_ and TRN_B_➔TC_C_ were both maximally weighted would be equally capable of supporting robust signal propagation, regardless of whether these synapses were varied homogeneously or heterogeneously. By contrast, top-scoring heterogeneous network variants better supported both oscillation [*t*(38)=13.88, *p*<0.0001] and optimization [*t*(38)=18.04, *p*<0.0001] than their homogeneous counterparts. Because networks supporting the propagation of oscillatory activity would, by definition, score high with respect to optimization, these results not only confirmed that heterogeneous networks were more likely than homogenous networks to accommodate this oscillatory mechanism, but furthermore that propagation of oscillation across the thalamocortical network was associated with higher oscillation scores than post-inhibitory-driven oscillation in TC_C_, the predominant form of oscillation observed in homogeneous networks.

**Figure 4.**
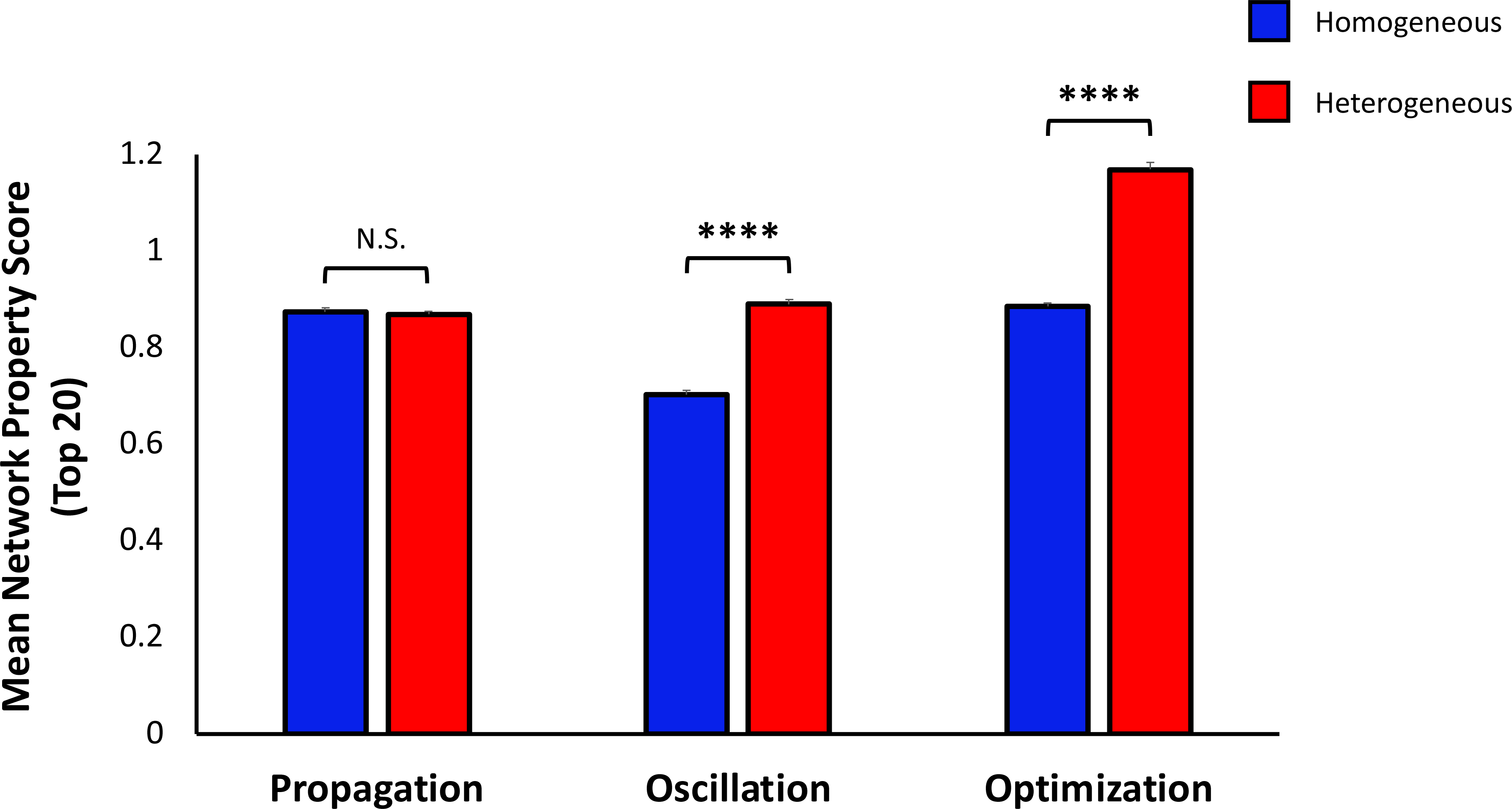
Propagation, as measured in those network permutations scoring highest with respect to the property, was equally supported in networks where synaptic weights varied independently of one another (heterogeneously; red) as in networks where synaptic strength varied homogeneously (blue) by class [unpaired *t*-test, *t*(38)=0.46, *p*=0.647]. By contrast, oscillation and optimization scores were significantly higher in top-performing heterogeneous networks than their homogeneous counterparts [oscillation: *t*(38)=13.88, *p*<0.0001; optimization: *t*(38)=18.04, *p*<0.0001]. Each bar corresponds to a mean of the top 20 network propagation or oscillation scores within each synaptic architecture group; error bars indicate standard errors of the mean. ****=*p*<0.0001; N.S.=not significant.

## Discussion

The simulations presented here suggest that open-loop TC-TRN-TC synaptic motifs (Fig. 1B, right) could function as a substrate for signal propagation across cortical networks without the need for direct cortico-cortical, intra-reticular or corticothalamic connectivity. Post-inhibitory rebound mediated by T-type calcium channels served as a substrate for both propagation and oscillation in the simulated networks. TRN-TRN connections, either chemical or electrical (Fig. 1B, left and middle), diminished horizontal propagation by disrupting the precise timing relationships required to propagate a signal across the network. Models with heterogeneously varied TRN synapses outperformed those whose synapses varied as a class with respect to the propagation of oscillatory activity, consistent with the emerging literature documenting cellular and synaptic heterogeneity in the TRN (70–72). These data suggest that widespread propagating cortical activity, under both pathological and physiological conditions, may be mediated, at least in part, by intrathalamic connections. The model makes strong predictions that can be tested physiologically. Finally, the approach used here, which employed supercomputing applications to search through a very large parameter space, serves as model for future computational models with large parameter spaces.

Like most of the thalamic (19, 47–50) and thalamocortical models (52, 53, 55) that inspired our model, we utilized single-compartment, Hodgkin-Huxley neurons. While these model cells contribute to the computational parsimony and practicality of network models, particularly where the analysis of network dynamics is prioritized, they neglect the intrinsic cable properties of real neurons and, relatedly, the spatially disparate nature of synaptic integration and heterogeneous expression of intrinsic and synaptic conductances (73, 74). Such considerations are particularly relevant here relative to dendritic distributions of T- and H-currents in TC neurons (52, 54, 75, 76) and TRN neurons (54, 77–79). Although multicompartment neuronal models incorporating such details could conceivably alter the network dynamics being studied, they were not necessary to simulate the propagation of oscillatory waves seen physiologically (18, 19, 48–50).

Additionally, the present model omitted explicit corticothalamic and corticoreticular synapses, both of which have been identified and physiologically characterized to varying degrees (80–86), though the former were effectively amalgamated with both feedforward sensory and modulatory projections to the thalamus in the form of the Poisson-modulated external input we delivered to individual TC neurons. Both forms of feedback have been implicated in the spread of spindle waves and in the maintenance of their synchronization over large distance scales (on the order of the length of the mammalian forebrain) and are furthermore known to drive spindle wave formation and propagation *in vivo* by polysynaptically recruiting TC neurons via TRN-mediated post-inhibitory rebound (80, 83, 86–91). It should be noted, however, that short-range coherence of spindle waves, which can be elicited in isolated thalamic slice preparations (18, 68), is preserved following decortication, both *in vivo* and *in silico* (52, 83, 88). By extension, it is reasonable to assume that the dynamics of the spindle-like waveforms generated in our small-scale, broadly feedforward model, in which the cortex served solely as an output layer, would not be qualitatively altered by corticothalamic or corticoreticular feedback.

### Comparison to related computational models and physiological data

Although the production of spindle waves was not an explicit objective of our study, some of the wave dynamics arising in our networks were nevertheless consistent with those inherent to spindle or spindle-like waves. Despite possessing higher degrees of TC➔TRN and TRN➔TC synaptic divergence and lacking the exclusively open-loop TC-TRN-TC architecture characterizing a subset of our network variants, other isolated thalamic models allowing for longitudinal wave propagation similarly accommodated this propagation along the lattice of interconnected TC and TRN neurons by way of laterally inhibitory TRN-TC synapses (19, 48, 50, 92); at short ranges, this mechanism of signal propagation also prevailed in larger-scale thalamo-reticulo-cortical models, while corticothalamic projections acted to propagate activity to more distal sites [(52); see (93), for a schematic illustrating short- and long-range thalamocortical wave propagation]. Comparably, recurrently inhibitory TRN-TC synapses have been documented to play a vital role in the generation of oscillatory behavior in the thalamus (17, 93). The temporal parameters of propagating and oscillation signals in our model also matched some of those previously reported: the mean signal propagation velocity and oscillation frequency measured across homogeneous networks fell within the ranges of spindle wave propagation velocities and intraspindle spike frequencies reported in both physiological and computational spindle wave studies (19, 48, 68, 94, 95).

Several key structural elements of our set of network models and the range of phenomena they produced distinguish them from previous thalamic and thalamocortical models. One particularly notable point of departure relative to similar network models was the extent to which thalamoreticular, reticulothalamic, and thalamocortical synapses diverged. Although all three classes of synapses are known to diverge significantly and have been observed to target neuronal somata hundreds of microns from their origins (25, 32, 58, 96–100), the TC-TRN, TRN-TC, and TC-L4 synapses in our model were constrained to remain strictly local and minimally divergent (or non-divergent, in the case of TC-TRN and TC-L4 synapses). With respect to the first two classes of synapses, this constraint was imposed to probe the impact the disynaptic TC-TRN-TC open-loop motifs characterizing a subset of network permutations, which constituted one of the foci of our study, and analyze the signal propagation they may support. This neuroanatomical scheme contrasted with previous computational models featuring parallel, interconnected thalamoreticular pathways, in which both TC and TRN synapsed bidirectionally with several neighboring TRN and TC cells, respectively, within a radius of several hundred microns (e.g., 19, 48–50, 52–54, 67, 101). It is highly likely that if more divergent synaptic connections were used in the current model, even greater propagation would have been observed.

### The functional implications open-loop thalamo-reticulo-thalamic synaptic motifs

The spread of activity from one cortical region to another is a foundational concept at the core of our understanding of sensory processing, higher order-cognitive functions such as attention and language, sleep-related oscillatory phenomena, and pathological findings such as propagation of ictal discharges and migraine. Despite the importance of communication between cortical regions, its underlying substrates are not well understood. It has long been speculated that the TRN could serve as a control point for large-scale cortical signal processing given its central location, the high degree of convergence of projections involved in attention, arousal and emotion onto the TRN and the TRN’s particularly strong control over TC firing properties (11–13, 102–104). Although the anatomical bases of open-loop TC-TRN-TC motifs have been partially characterized, their functional significance in the brain lingers as a subject of continued speculation. Here we show that open-loop TC-TRN-TC architectures can support at least short-range cortical signal propagation. Within the thalamus, these configurations have thus far been observed both within and across individual thalamic nuclei and are thought to serve as pathways for intra- and cross-modal modulation, respectively (32, 34–40); as has been previously surmised, these synaptic pathways could also plausibly lend themselves to sensory enhancement, multisensory integration, and attentional mechanisms (10, 35, 46, 105). At a minimum, and as inferred from physiological studies, open-loop pathways should be fully capable of supporting signaling propagation from one thalamic relay neuron to another through a limited number of intervening synapses (with a disynaptic pathway serving as the shortest such configuration). Moreover, interference with thalamoreticular transmission should cause a breakdown in some forms of cortical signal propagation. Recent work has established that stimulation of the TRN *in vivo* can induce propagating rhythmic activity across the cortex (106–108). These data suggest that abnormal cortical signal propagation seen in seizures, migraines or hallucinations may be disrupted by targeted therapeutics applied to the TRN. Both forthcoming physiological investigation and future modeling studies will be able to evaluate these predictions and help provide a full accounting of the role of the various modes of connectivity between cortical regions.

## Methods

### Intrinsic neuronal models

Our network model was directly based on an earlier incarnation published by our research group (46). Single-compartment TC, TRN, and L4 model neurons obeyed Hodgkin-Huxley kinetics, with membrane potentials *V* varying according to the first-order differential equation

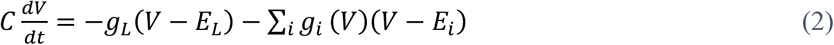

where *C* is the membrane capacitance, *g*_*L*_ and *E*_*L*_ are the leakage conductance and reversal potential, respectively, and *g*_*i*_*(V)* and *E*_*i*_ are the dynamic conductance and reversal potential, respectively, of the *i*th voltage-gated, ligand-gated (chemical synaptic), or electrical synaptic conductance (for electrical synaptic conductances, the effective reversal potential is equal to the presynaptic membrane potential; see Equation 3a). All three varieties of model neurons expressed both the standard transient sodium (*I*_*Na*_) and delayed-rectifier potassium (*I*_*K*_) currents, as reported by (46). TC and TRN neurons additionally included a T-type calcium conductance (T-current; *I*_*T*_) and hyperpolarization-activated cation current (H-current; *I*_*H*_), following the TC model of (109). Both TRN and L4 cells expressed a slow, non-inactivating potassium conductance (*I*_*M*_), following the modeling of (110), which accounts for the spike-frequency adaptation previously reported in physiological recordings from these neurons (46, 111). A list of intrinsic model cell parameters, including current conductances, reversal potentials, selected gating kinetics, and membrane capacitance, can be found in Table S3.

### Synaptic models

The kinetics of chemical synapses in our model network conformed to the synaptic depression model of (112), following our previous computational network model (46). This model presupposes a finite quantity of “resources,” akin to synaptic vesicles, capable of being released by the presynaptic neuron; these resources can exist in an active, inactive, or recovered state. A parameter *U*_*SE*_ characterizes the fraction of recovered resources that can be converted to an active state (i.e., for release by the presynaptic neuron) following action potential induction in the presynaptic axon terminal(s). Following resource activation, synapses inactivate according to the time constant *τ*_inact_; resources become available again for activation after a recovery period described by the time constant *τ*_recov_. These parameters, along with the neurotransmitters, postsynaptic conductances, and reversal potentials characterizing all of the chemical synapses in our model, are given in Table S4.

Glutamatergic thalamoreticular and thalamocortical (TC-L4) and baseline GABAergic reticulothalamic synaptic parameters matched those of our earlier model (46), with the latter synapses allowed to vary in conductance. TRN-TC signaling was mediated exclusively through GABA_A_ receptors, mirroring other thalamic and thalamocortical models in which the slower TRN-TC GABA_B_ conductance was omitted (51, 54, 55). Both GABAergic (TRN-TRN_GABA_) and electrical synapses (TRN-TRN_Elec_) were included between TRN neurons; as with TRN-TC synapses, both varieties of TRN-TRN synapses were allowed to vary in strength. Although evidence has been presented challenging the existence of GABAergic intrareticular synapses in certain mammalian species and age groups (10, 63, 113–115), our model avoided making assumptions regarding their presence, strength, or spatial distribution by allowing the associated synaptic conductances to vary over a range of physiological values, including zero, and in distribution. The reversal potential, conductance, and kinetics of the external synapses projecting to the TC neurons were directly based on retinogeniculate synapses (116), although the generic nature of the external inputs in our model allows them to represent not only immediately upstream sensory input but also brainstem modulation (e.g., serotonergic, adrenergic) known to act on thalamic nuclei (117).

Electrical synapses between TRN neurons were based on the Cx36-dependent intrareticular gap junctions first identified by (58). For TRN neurons, the sum of electrical synaptic currents (*I*_*Elec*_) entering any postsynaptic neuron *j* from presynaptic neurons *i* was included in the rightmost term from Equation 1 and calculated as

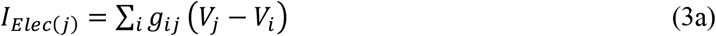

where *g*_*ij*_ was itself calculated as

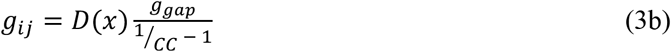

where *CC* was the electrical coupling coefficient between TRN neurons *i* and *j*, *g*_*gap*_ is the gap junction conductance (set at 5 nS), and *D(x)* was a scaling factor that depended on the physical distance between the coupled TRN neurons (54, 73, 118). TRN-TRN_Elec_ synapses were symmetrical (non-rectifying), such that *G*_*ij*_=*G*_*ji*_.

We extrapolated the attenuation of intrareticular synaptic strength as a function of intracellular distance based on mappings of intrinsic connections within the TRN along a horizontal (anteroposterior) plane assembled by (61). Assuming 1) an intracellular distance of 50 µm between adjacent TRN neurons, 2) a distance *x* (in multiples of 50 µm) between non-adjacent neurons, and 3) a Gaussian falloff in synaptic strength (119), the baseline (adjacent-neuron) conductances of TRN-TRN_GABA_ and TRN-TRN_Elec_ synapses were scaled for non-adjacent synapses using the function

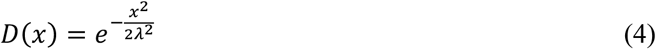

where λ_GABA_=531 µm and λ_Elec_=130 µm.

Given the small spatial scale of our model, synaptic delays associated with finite axonal conductance times within the TRN and between the TRN and dorsal thalamus were disregarded, mirroring the simplification incorporated into previous thalamic and thalamocortical models simulating synaptic interactions on the order of 100 microns (48, 54). Although small (~1 ms) thalamocortical delays were inserted into the network model of (54), these were likewise omitted on the basis of the cortex functioning solely as an output layer in our model.

### Computations and statistics

Our model was coded, simulated, and analyzed in MATLAB R2018b (MathWorks), utilizing both a Dell Inspiron 3847 and Hewlett-Packard Z840 running Windows 10 and nodes on the Illinois Campus Cluster (National Center for Supercomputing Applications, University of Illinois at Urbana-Champaign). Simulations employed 0.1-ms time steps, with temporal integration based on the hybrid analytic-numeral integration method of (120), which optimizes between accurate solutions to Hodgkin-Huxley and synaptic models and computational efficiency. All simulations commenced with a 200-ms equilibration period, during which no external stimulation was delivered to TC neurons; this allowed all network elements to attain steady-state conditions. Statistical analysis was performed in both MATLAB and R (121), with the *glmnet* package (122) utilized within the latter platform to perform regression analyses. Multiple linear regression was employed to establish rudimentary relationships between synaptic classes (homogeneously synaptic networks) or individual synapses (heterogeneously synaptic networks) and each of the two studied network properties, even in instances where these relationships deviated from linearity. 2° regression models with interaction terms elucidated how synaptic interactions and nonlinearities affected these network properties. Regressions were optimized using elastic net regularization, with the specific regularization hyperparameter α selected to minimize each regression model’s root-mean-square error. To convey the relative influence of different synaptic classes or individual synapses on dynamic network properties, all regression coefficients are reported here as normalized to the coefficient with the largest absolute value; the effects corresponding to NRCs with absolute values of less than 0.05 were disregarded as negligibly influential on network dynamics. Both unpaired Student *t*-tests and one-way ANOVA models were used to compare the mean property scores between different sets of networks, with Tukey’s honestly significant difference tests used to ascertain pairwise difference between groups in the latter. Kolmogorov-Smirnov and Levene’s tests were employed to confirm normality and homogeneity of variance, respectively, when utilizing parametric mean-comparison tests; data were log-transformed as needed to conform to these prerequisites.

## Acknowledgments

This work made use of the Illinois Campus Cluster, a computing resource that is operated by the Illinois Campus Cluster Program (ICCP) in conjunction with the National Center for Supercomputing Applications (NCSA) and which is supported by funds from the University of Illinois at Urbana-Champaign. We additionally thank Profs. Thomas Anastasio, Ruoqing Zhu, and Rama Ratnam, Drs. Kush Paul and Baher Ibrahim, and Mr. Weddie Jackson for their valuable analytical, statistical, and technical insights.

---

**Table S1.** Normalized linear and second-order regression coefficients for propagation and oscillation in homogeneously varied synaptic networks.

| Normalized Regression Coefficients (NRCs) |  |  |  |  |
| --- | --- | --- | --- | --- |
| Synaptic Variable | Propagation Linear | Propagation 2° | Oscillation Linear | Oscillation 2° |
| TRN-TRN<sub>GABA</sub> | -0.173 | -0.670 | - | 0.060 |
| TRN-TRN<sub>Elec</sub> | -0.136 | -0.347 | - | - |
| TRN-TC | 1.000 | 1.000 | -1.000 | -0.052 |
| (TRN-TRN<sub>GABA</sub>)<sup>2</sup> | - | 0.332 | - | - |
| (TRN-TRN<sub>Elec</sub>)<sup>2</sup> | - | 0.164 | - | - |
| (TRN-TC)<sup>2</sup> | - | 0.594 | - | -1.000 |
| TRN-TRN<sub>GABA</sub> x TRN-TRN<sub>Elec</sub> | - | 0.262 | - | - |
| TRN-TRN<sub>GABA</sub> x TRN-TC | - | -0.152 | - | - |
| TRN-TRN<sub>Elec</sub> x TRN-TC | - | -0.365 | - | - |

• The regressions include 1°, 2°, and interaction terms corresponding to TRN-TRN<sub>GABA</sub>, TRN-TRN<sub>Elec</sub>, and open-loop TC-TRN-TC synapses/pathways. Terms associated with regression coefficients of absolute values < 0.05 are omitted. Positive and negative terms are highlighted in red and blue, respectively. Linear regression for propagation, $R^2=0.793$ , RMSE=0.047, $p<0.0001$ ; second-order regression for propagation, $R^2=0.842$ , RMSE=0.041, $p<0.0001$ ; linear regression for oscillation, $R^2=0.526$ , RMSE=0.145, $p<0.0001$ ; second-order regression for oscillation, $R^2=0.630$ , RMSE=0.128, $p<0.0001$ .

**Table S2.**
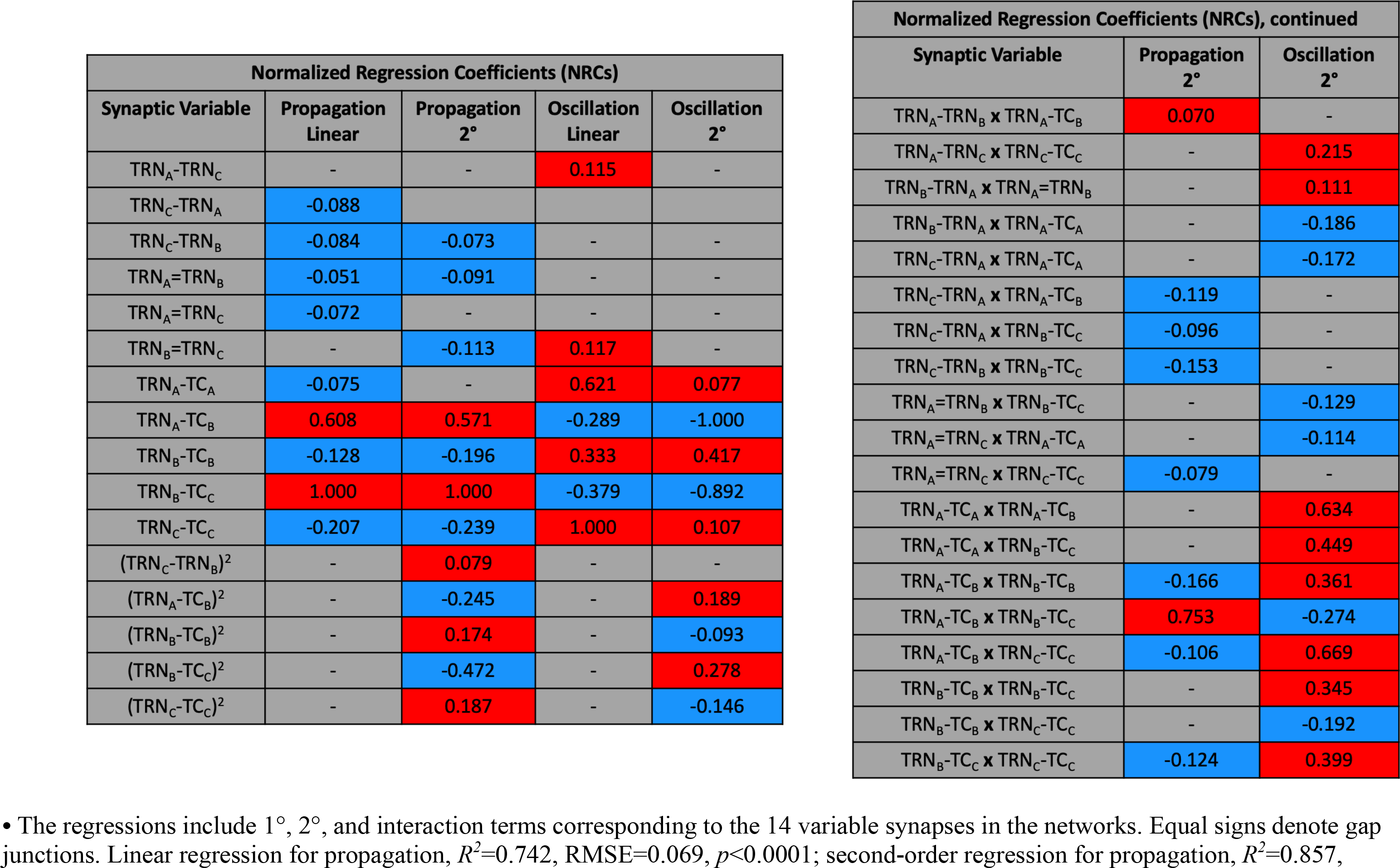
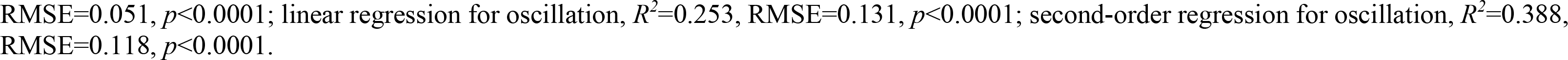
Normalized linear and second-order regression coefficients for propagation and oscillation in heterogeneously varied synaptic networks.

**Table S3.** Intrinsic model cellular parameters.

| Model Cellular Parameters |  |  |  |
| --- | --- | --- | --- |
| Parameter | TC cell | TRN cell | L4 cell |
| Leak conductance, $g_L$ (nS) | 3.263 | 3.7928 | 4.8128 |
| Leak reversal potential, $E_L$ (mV) | -60.03 | -57 | -60.2354 |
| Transient sodium conductance, $g_{Na}$ (nS) | 1,500 | 3,000 | 3,000 |
| Sodium equilibrium potential, $E_{Na}$ (mV) | 50 | | |
| Delayed-rectifier potassium conductance, $g_K$ (nS) | 520 | 400 | 140 |
| M-type potassium conductance, $g_M$ (nS) | - | 3.5 | 1.5 |
| M-type potassium time constant, $\tau_M$ (ms) | - | 200 | 180 |
| Potassium equilibrium potential, $E_K$ (mV) | -100 | | -90 |
| T-type calcium conductance, $g_T$ (nS) | 45 | 21 | - |
| Calcium equilibrium potential, $E_T$ (mV) | 120 | | |
| H-current conductance, $g_H$ (nS) | 0.608 | 0.0192 | - |
| H-current reversal potential, $E_H$ (mV) | -33 | | - |
| Membrane capacitance, $C_m$ (pF) | 100.4 | 75.0 | 109.3865 |

**Table S4.** Model synaptic parameters.

| Model Synaptic Parameters |  |  |  |  |  |  |
| --- | --- | --- | --- | --- | --- | --- |
| Synapse | Neurotransmitter | Conductance (nS) | $\tau_{\text{recov}}$ (ms) | $\tau_{\text{inact}}$ (ms) | Reversal Potential (mV) | $U_{SE}$ |
| External synapse to TC cell | (Glutamate) | 32 | 125 | 2.64 | 0 | 0.76 |
| TC-to-TRN cell synapse (TC-TRN) | Glutamate | 150 | 500 | 2.64 | 0 | 0.76 |
| TC-to-L4 cell synapse (TC-L4) | Glutamate | 50 | 160 | 11.52 | 0 | 0.8113 |
| TRN-to-TC cell synapse (TRN-TC) | GABA<sub>A</sub> | Variable (0-80) | 167.29 | 16.62 | -80 | 0.62 |
| Chemical TRN-to-TRN cell synapse (TRN-TRN<sub>GABA</sub>) | GABA<sub>A</sub> | Variable (0-450) | 225 | 15 | -75 | 0.62 |

